# Common roles for serotonin in rats and humans for computations underlying flexible decision-making

**DOI:** 10.1101/2023.02.15.527569

**Authors:** Qiang Luo, Jonathan W. Kanen, Andrea Bari, Nikolina Skandali, Christelle Langley, Gitte Moos Knudsen, Johan Alsiö, Benjamin U. Phillips, Barbara J. Sahakian, Rudolf N. Cardinal, Trevor W. Robbins

## Abstract

Serotonin is critical for adapting behavior flexibly to meet changing environmental demands. Cognitive flexibility is important both for successful attainment of goals, as well as for social interactions, and is frequently impaired in neuropsychiatric disorders, including obsessive-compulsive disorder (OCD). However, a unifying mechanistic framework accounting for the role of serotonin in behavioral flexibility has remained elusive. Here, we demonstrate common effects of manipulating serotonin function across two species (rats and humans) on latent processes supporting choice behavior during probabilistic reversal learning using computational modelling. The findings support a role of serotonin in behavioral flexibility and plasticity, indicated, respectively, by increases or decreases in choice repetition (‘stickiness’) or reinforcement learning rates depending upon manipulations intended to increase or decrease serotonin function. More specifically, the rate at which expected value increased following reward and decreased following punishment (reward and punishment ‘learning rates’) was greatest after sub-chronic administration of the selective serotonin reuptake (SSRI) citalopram (5 mg/kg for 7 days followed by 10 mg/kg twice a day for 5 days) in rats. Conversely, humans given a single dose of an SSRI (20mg escitalopram), which can decrease post-synaptic serotonin signalling, and rats that received the neurotoxin 5,7-dihydroxytryptamine (5,7-DHT), which destroys forebrain serotonergic neurons, exhibited decreased reward learning rates. A basic perseverative tendency (‘stickiness’), or choice repetition irrespective of the outcome produced, was likewise increased in rats after the 12-day SSRI regimen and decreased after single dose SSRI in humans and 5,7-DHT in rats. These common effects of serotonergic manipulations on rats and humans – identified via computational modelling – suggest an evolutionarily conserved role for serotonin in plasticity and behavioral flexibility and have clinical relevance transdiagnostically for neuropsychiatric disorders.

## Introduction

Humans and other animals alike must maximise rewards and minimise punishments to survive and thrive. Across phylogeny this involves learning about cues or locations that inform whether an action is likely to result in a good or bad outcome. Adaptive behavior, however, must also be flexible: the ability to disengage from previously learned actions that are no longer useful or appropriate to the situation is fundamental to well-being. Indeed, behavior can become abnormally stimulus-bound and perseverative in compulsive disorders ^1-5^. Furthermore, learning the best course of action can require withstanding occasional negative feedback, which should sometimes be ignored if rare. Indeed, inappropriately switching behavior away from an adaptive action following misleading or even negative feedback (‘lose-shift’) has been reported across several traditional psychiatric diagnostic categories ^6-10^.

The neurotransmitter serotonin (5-hydroxytryptamine; 5-HT) is widely implicated in behavioral flexibility^11-18^. Perturbing 5-HT function can affect both perseveration and lose-shift behavior, which are commonly assessed using probabilistic reversal learning (PRL) paradigms (Figure 1 A-B): a subject learns through trial and error the most adaptive action in a choice procedure, the contingencies of which eventually reverse, sometimes repeatedly ^12, 19-21^. A unifying framework for 5-HT in these processes has, however, remained elusive. To this end, we proposed to use a mechanistic modelling framework to align behavioral changes in PRL following serotonergic manipulations in rats ^19^ and humans ^22^.

**Figure 1.**
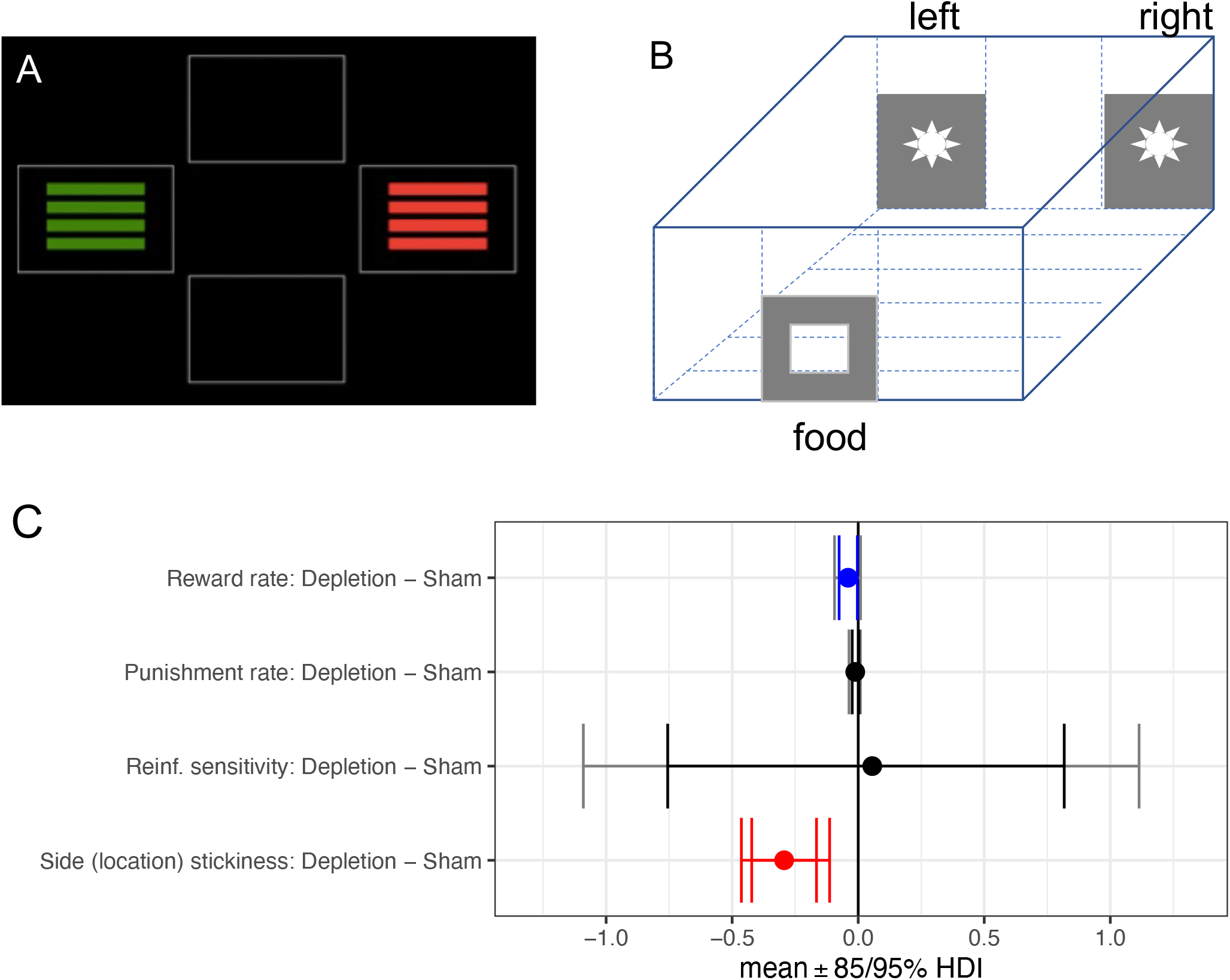
Task schematics for probabilistic reversal learning and effects of serotonin depletion on model parameters in rats. **A)** Experiment in humans (example trial on touchscreen computer) and **B)** Experiment in rats (two apertures illuminated simultaneously to the left and right of a central aperture with reinforcement contingencies 80% : 20% for left : right or right : left, and a food pellet was given to a food magazine positioned on the opposite wall of the operant chamber if the rewarding location was chosen). **C**) Side (location) stickiness was diminished by neurotoxic 5-HT depletion, *i*.*e*., 5,7-dihydroxytryptamine. Reinf. = reinforcement. Red signifies a difference between the parameter per-condition mean according to the Bayesian “credible interval”, 0 ∉ 95% HDI. Blue signifies a significance by the 85% HDI. The inner interval represents the 85% HDI, while the outer interval represents the 95% HDI.

Reinforcement learning (RL) is a well-established computational mechanism for the analysis of latent mechanisms underlying choice behavior as it unfolds dynamically over time ^23^. Standard RL models typically conceptualise choice in relation to an action’s value, derived from an accumulated reinforcement history, and incorporate parameters that estimate how quickly action values are learned (‘learning rate’) and the extent to which that value is acted upon (often termed ‘inverse temperature’ in relation to the mathematical softmax function typically used; here, termed ‘reinforcement sensitivity’) ^24^. Stickiness parameters, by contrast, track the extent to which behavioral tendencies are shaped by engagement with discrete cues (stimuli) or locations, irrespective of an action’s outcome. Stickiness can therefore be considered a value-free component of behavior ^25, 26^. Across six previously published experiments in rats and humans and a recently published computational modelling study in humans, we examined whether stickiness or other RL parameters (learning rates or reinforcement sensitivity) contributed meaningfully to behavior, and examined whether 5-HT function would consistently modulate any of these parameters across species.

The stickiness parameter has recently emerged as important for understanding compulsivity: stickiness was significantly high in stimulant use disorder (SUD) but abnormally low in obsessive–compulsive disorder (OCD) during PRL performance ^7^. Meanwhile, value-free influences have been notably absent from prominent computational accounts of goal-directed (or ‘model-based’) versus habitual (or ‘model-free’) controllers of behavior ^26^. These have traditionally revolved around environmental features relevant to outcomes ^27, 28^. This has hindered contextualisation within the rich literature on the neural basis of habits (reinforcer-independent perseveration) ^29^. A traditional view of stimulus–response habits is that they are created and strengthened by reinforcement, acting to enhance direct links between environmental stimuli and responses ^30^; they are thus “model-free” in that they do not involve representations of the expected consequences of behavior, but are “value-based” in that they are created by valenced reinforcement. However, there are other aspects of behavior that are independent of reinforcement or value. Indeed, value-free (action outcome-irrelevant) factors similar to stickiness were recently shown to be important for understanding goal-directed decision-making ^28^. Accounting for stickiness – value-free perseveration – may therefore aid in better dissecting the nature of imbalanced goal-directed versus habitual behavior seen in OCD, SUD, and other conditions ^31-33^, a balance that is sensitive to serotonergic disruption in humans and rodents ^34-36^.

Two common methods for studying serotonin are through serotonin depletion and treatment with selective serotonin reuptake inhibitors (SSRIs). In non-human animals, depletion can be achieved using the neurotoxin 5,7-dihydroxytryptamine (5,7-DHT) which produces a profound loss of serotonergic fibers ^37^. SSRIs, meanwhile, are first-line pharmacological treatments for several psychiatric conditions including major depressive disorder (MDD) ^38^, anxiety disorders ^39^, post-traumatic stress disorder (PTSD) ^40^, and OCD ^41^, yet both the computational and neural mechanisms underlying their efficacy remain poorly understood. SSRIs block the 5-HT transporter and thus reuptake of 5-HT, which increases extracellular serotonin levels; however, this occurs not only in projection areas but also in the vicinity of 5-HT_1A_ somatodendritic autoreceptors, activation of which leads to decreased firing rates of 5-HT neurons ^42^. SSRIs can thus paradoxically lower 5-HT concentrations in projection regions when given acutely, especially at low doses ^43^, and firing rates return to baseline after 5-HT_1A_ autoreceptors are desensitised by repeated administration ^44^. This mechanism might be reflected in a delayed clinical onset of the treatment effect of SSRI on mood ^45^. For this reason, effects of both acute and chronic SSRIs in rats were studied, with the prediction that a higher acute dose and a chronic use could overcome these feedback effects of a low acute dose and produce an increase in serotonin transmission ^19^. The 20mg used in the acute study with healthy humans ^22^, while within the therapeutic range, is a lower acute dose than used in some experimental animal studies.

Here, the primary question was whether serotonergic manipulations would cause similar perturbations of model parameters across both rats and humans, thereby demonstrating the evolutionary significance of the role of serotonin in cognitive flexibility. As an increased tendency for lose-shift behavior induced by acute SSRI has been conceptualised as hypersensitivity to negative feedback ^19, 22^, we asked whether this would be reflected in elevated punishment learning rates. Selective 5-HT depletion via 5,7-DHT of the orbitofrontal cortex (OFC) or amygdala in marmoset monkeys, meanwhile, reduced reinforcement learning rates (for rewards or punishments), and modulated stickiness ^46^; we hypothesised that changes in learning rate or stickiness parameters would occur following global 5-HT manipulations in rats and humans. We predicted that incorporating stickiness parameters would be central to capturing effects of 5-HT on behavioral flexibility and would increase or decrease depending on changes in serotonin transmission.

## Materials and Methods

### Probabilistic reversal learning task: humans

The task used in the human SSRI experiment ^22^ is shown in Figure 1A, and contained 80 trials: 40 during acquisition and 40 following reversal. In other words, there was a fixed number of trials and a single reversal. For the first 40 trials, one option yielded positive feedback on 80% of trials, the other option on 20% of trials. These contingencies reversed for the latter 40 trials. Positive feedback was given in the form of the word “CORRECT” on the touchscreen computer and a high tone, negative feedback was conveyed by the word “WRONG” and a low tone. The task was self-paced.

### Probabilistic reversal learning task: rats

Following training and determination of stable levels of accuracy and a lack of side bias ^19^ in operant chambers controlled by the Whisker control system ^47^, rats were presented with two apertures illuminated simultaneously to the left and right of a central (inactive) aperture (Figure 1B). Responding at the ‘correct’ location was associated with an 80% probability food reward (and 20% probability of a time-out punishment), whereas responding at the ‘incorrect’ location yielded reward on only 20% of trials (and punishment on 80%). Reward was in the form of a 45 mg food pellet (Noyes dustless pellets; Sandown Scientific, Middlesex, UK) delivered to a food magazine positioned on the opposite wall of the operant chamber. Punishment was given in the form of a 2.5-second time-out. The left and right apertures were illuminated for 30 seconds signifying the response window. The next trial was triggered by retrieval of the pellet from the magazine. If no response was made, the trial was categorised as an omission and resulted in a 5-second time-out. Responding to an unlit aperture had no programmed consequence. Reversals occurred after the animal made eight consecutive correct responses, at which point the correct aperture became the incorrect aperture and vice versa. A session consisted of 200 trials to be completed during a 40-minute period. One session was conducted per day.

### 5,7-DHT forebrain 5-HT depletion: rats

Sixteen rats were included in the final analysis. Rats were pre-treated intraperitoneally (i.p.) with 20 mg/kg of desipramine hydrochloride (Sigma, Poole, UK) in order to preserve noradrenergic neurons. Half of the rats were randomly assigned to receive bilateral intracerebroventricular (i.c.v.) infusions of 80 μg 5,7-DHT creatinine sulfate diluted in 10 μg of 10% ascorbic acid in saline, guided by a stereotaxic frame, whilst the other half received a sham infusion of 10 μg 0.01 M phosphate-buffered saline (PBS) – vehicle ^19^. Post-mortem neurochemistry confirmed that 5,7-DHT infusions produced a near-total depletion of brain serotonin and decreased levels of the serotonin metabolite 5-hydroxyindoleacetic acid (5-HIAA) relative to controls in all regions examined: OFC, prelimbic cortex, anterior cingulate cortex, nucleus accumbens, dorsomedial striatum, dorsolateral striatum, amygdala, dorsal hippocampus (all p<.05)^19^. Levels of dopamine, norepinephrine, and the dopamine metabolite dihydroxyphenylacetic acid (DOPAC) were not significantly different from controls in any of these regions (all p > .05)^19^. Data were analysed from seven consecutive sessions conducted following surgery in the previous report ^19^. Computational model convergence was achieved when modelling behavior from all seven sessions collectively, which is reported in the current study. Conversely, computational model convergence could not be achieved when modelling the seven sessions separately.

### SSRI administration: rats

Animals were divided into groups matched for task accuracy and then randomly assigned via a Latin square design to receive injections i.p. of either citalopram hydrobromide (1 mg/kg or 10 mg/kg; Tocris, Bristol, UK). Citalopram, dissolved in 0.01 M PBS, or vehicle was administered 30 minutes before the task ^19^. Eleven rats were included in the final analysis after receiving vehicle, 1 mg/kg, or 10 mg/kg citalopram ^19^. Fourteen rats were included in the repeated and sub-chronic citalopram experiment. The citalopram group was administered 5 mg/kg citalopram 30 min before testing, for seven consecutive days (*n*=7). The vehicle group (*n*=7), instead, received the same number of daily injections of 0.01 M phosphate-buffered saline ^19^. After seven days, the citalopram group received 10 mg/kg of citalopram twice a day (about 4 h before the testing) for five consecutive days, to study the long-lasting effects of sub-chronic dosing ^19^.

All the above animal experiments were conducted in accordance with the United Kingdom Animals (Scientific Procedures) Act, 1986 (PPL 80/2234) in our previous study ^19^.

### SSRI administration: humans

The protocol was ethically approved (Cambridge Central NHS Research Ethics Committee, reference 15/EE/0004). Volunteers gave informed consent and were paid. Participants were healthy and without a personal or family history of psychiatric or neurological disorders ^22^. In a randomised, double-blind, placebo-controlled, between-groups design ^22^, healthy volunteers received either escitalopram (*n*=32) or placebo (*n*=33). The PRL task was conducted following a 3-hour waiting period after oral drug administration to attain peak plasma escitalopram concentration ^48^. Plasma analysis (*n*=59) verified increased escitalopram concentration ^22^ at 2.5 hours after the dose (t_54_ = 18.835, p < 0.001, mean = 14 ng/ml, standard deviation [SD] = 5.72) just before the task administration, and at 5.5 hours (t_54_ = 20.548, p < 0.001, mean = 17.24 ng/ml, SD = 4.27). Mood ratings were unaffected by single dose escitalopram administration (p > .05). There were no differences between groups in age, sex, years of education, depressive symptoms, or trait anxiety (all p > .05).

### Computational modelling of behavior

#### Overview

These methods are based on Kanen *et al*. ^7^. Four RL models were fitted to the behavioral data, which incorporated parameters that have been studied previously using a hierarchical Bayesian method^7, 49^. Models were fitted via Hamiltonian Markov chain Monte Carlo sampling implemented in Stan 2.17.2^50^. Convergence was checked according to 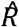, the potential scale reduction factor measure^51, 52^, which approaches 1 for perfect convergence. Values below 1.1 are typically used as a guideline for determining model convergence and 1.1 as a stringent criterion ^51^. In the current study, most of the models had an 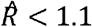, except for Model 4 in the sub-chronic 10 mg/kg experiment in rats 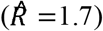 and Model 1 in the 5,7-DHT experiment in rats 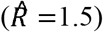. We assumed the four models examined had the same prior probability (0.25). Models were compared via a bridge sampling estimate of the likelihood ^53^, using the “bridgesampling” package in R ^54^. Bridge sampling directly estimates the marginal likelihood, and therefore the posterior probability of each model given the data (and prior model probabilities), under the assumption that the models represent the entire group of those to be considered. Posterior distributions were interpreted using the highest density interval (HDI) of posterior distributions, which is the Bayesian “credible interval”, at different significance levels including 75%, 80%, 85%, 90% and 95%. Together with the HDI, the group mean difference (MD) was also reported. The priors used for each parameter are shown in Supplemental Table 1. For the human experiments, trials were sequenced across all 80 trials of the PRL task, and on each trial the computational model was supplied with the participant’s identification number and condition, whether the trial resulted in positive or negative feedback, and which visual stimulus was selected. For the rat experiments, trials were sequenced across all sessions conducted under a given manipulation, and the computational model was supplied with the same information, but instead with the location of the aperture selected rather than the identification of the stimulus selected. Omissions were rare and they were not included in the computational analysis.

**Table 1.**
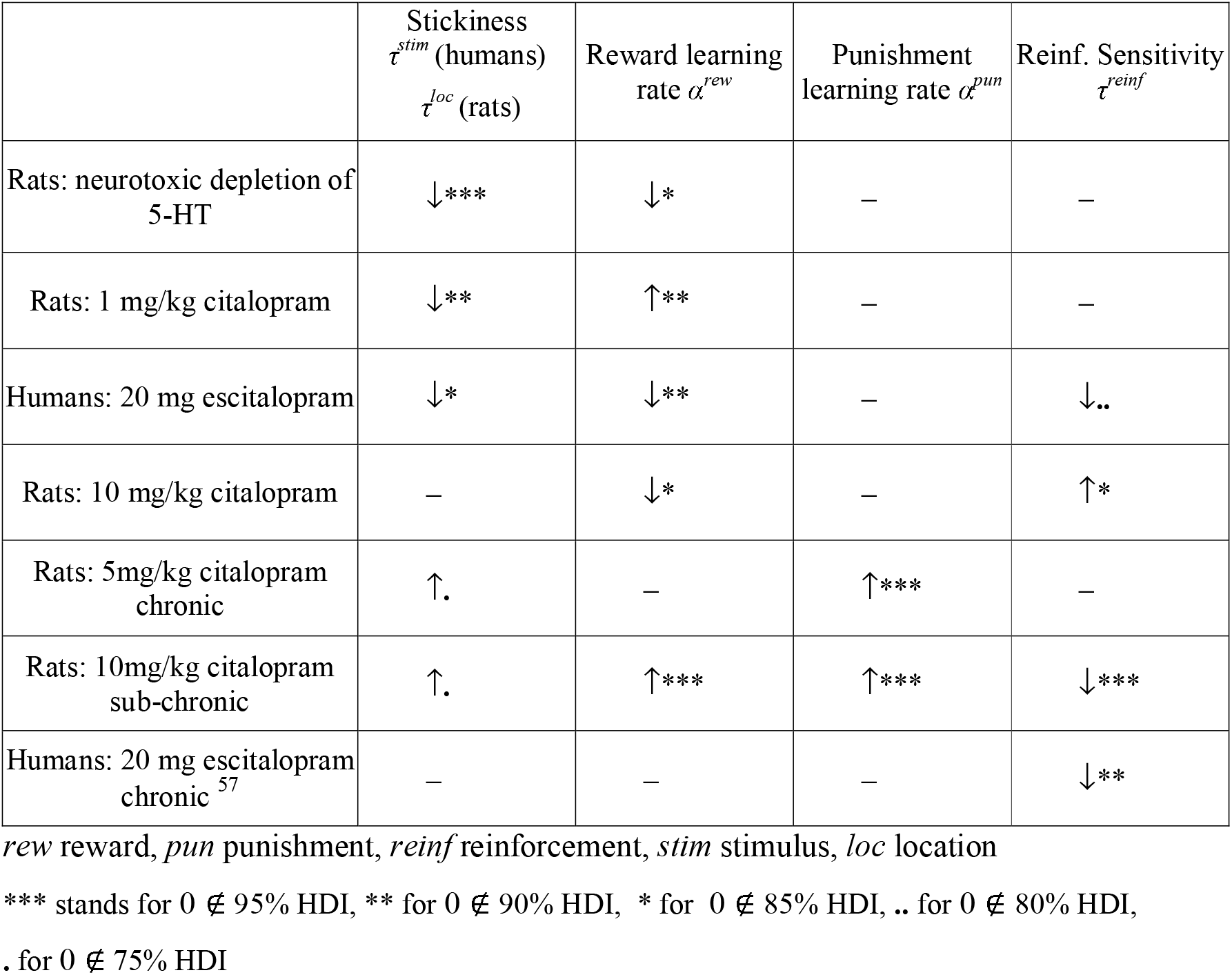
Summary of learning parameter effects.

| | Stickiness<br>$\tau^{stim}$ (humans)<br>$\tau^{loc}$ (rats) | Reward learning<br>rate $\alpha^{rew}$ | Punishment<br>learning rate $\alpha^{pun}$ | Reinf. Sensitivity<br>$\tau^{reinf}$ |
| --- | --- | --- | --- | --- |
| Rats: neurotoxic depletion of 5-HT | ↓*** | ↓* | — | — |
| Rats: 1 mg/kg citalopram | ↓** | ↑** | — | — |
| Humans: 20 mg escitalopram | ↓* | ↓** | — | ↓.. |
| Rats: 10 mg/kg citalopram | — | ↓* | — | ↑* |
| Rats: 5mg/kg citalopram chronic | ↑. | — | ↑*** | — |
| Rats: 10mg/kg citalopram sub-chronic | ↑. | ↑*** | ↑*** | ↓*** |
| Humans: 20 mg escitalopram chronic<sup>57</sup> | — | — | — | ↓** |

*rew* reward, *pun* punishment, *reinf* reinforcement, *stim* stimulus, *loc* location
\*\*\* stands for $0 \notin 95\%$ HDI, \*\* for $0 \notin 90\%$ HDI, \* for $0 \notin 85\%$ HDI, .. for $0 \notin 80\%$ HDI,
. for $0 \notin 75\%$ HDI

#### Models

Model 1 incorporated three parameters and was used to test the hypothesis that 5-HT would affect how positive versus negative feedback guides behavior. Separate learning rates for positive feedback (reward) α^*rew*^ and negative feedback (nonreward/punishment) α^*pun*^ were implemented. Positive reinforcement led to an increase in the value *V*_*i*_ of the stimulus *i* that was chosen, at a speed governed by the *reward learning rate* α^*rew*^, via *V*_*i,t*+1_ ← *V*_*i,t*_ + α^*rew*^(*R*_*t*_ – *V*_*i,t*_). *R*_*t*_ represents the outcome on trial *t* (defined as 1 on trials where positive feedback occurred), and (*R*_*t*_ – *V*_*i,t*_) the prediction error. On trials where negative feedback occurred *R*_*t*_ = 0, which led to a decrease in value of *V*_*i*_ at a speed governed by the *punishment learning rate* α^*pun*^, according to *V*_*i,t*+1_ ← *V*_*i,t*_ + α^*pun*^(*R*_*t*_ – *V*_*i,t*_). Stimulus value was incorporated into the final quantity controlling choice according to *Q*^*reinf*^_*t*_ = τ^*reinf*^*V*_*t*_. The additional parameter τ^*reinf*^, termed *reinforcement sensitivity*, governs the degree to which behavior is driven by reinforcement history. The quantities *Q* associated with the two available choices, for a given trial, were then input to a standard softmax choice function to compute the probability of each choice:

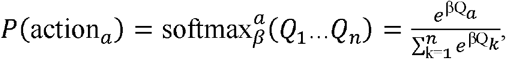

for *n*=2 choice options. The probability values for each trial emerging from the softmax function (*i*.*e*., the probability of choosing stimulus 1) were fitted to the subject’s actual choices (*i*.*e*., did the subject choose stimulus 1?). Softmax inverse temperature was set to β = 1, and as a result the reinforcement sensitivity parameter (τ^*reinf*^) directly represented the weight given to the exponents in the softmax function.

Model 2 was as model 1 but for the human experiments incorporated a “stimulus stickiness” parameter τ^*stim*^, which measures the tendency to repeat a response to a specific perceptual stimulus, irrespective of the action’s outcome. For the rat experiments a “side (location) stickiness” parameter τ^*loc*^ was substituted, which measures the tendency to repeat a response to a specific aperture in the operant chamber. Incorporating these two different stickiness parameters, depending on the species, accounts for task differences between the human and rat PRL experiments. This four-parameter model served to test whether accounting for stimulus-response learning, in addition to learning about action-outcome associations, would best characterise behavior. The stimulus stickiness effect was modelled as *Q*^*stim*^_*t*_ = τ^*stim*^*s*_*t–1*_, where *s*_*t–1*_ was 1 for a stimulus that was chosen on the previous trial and was otherwise 0. The final quantity controlling choice incorporated this additional parameter as *Q*_*t*_ = *Q*^*reinf*^_*t*_ + *Q*^*stim*^_*t*_. Quantities *Q*, corresponding to the two choice options on a given trial, were then fed into the softmax function as above.

Model 3 incorporated three parameters and served to test whether a single learning rate α^*reinf*^, rather than separate learning rates for rewards and punishments, optimally characterised behavior. Reward led to an increase in the value *V*_*i*_ of the stimulus *i* that was chosen, at a speed controlled by the *reinforcement rate* α^*reinf*^, via *V*_*i,t*+1_ ← *V*_*i,t*_ + α^*reinf*^(*R*_*t*_ – *V*_*i,t*_). *R*_*t*_ represents the outcome on trial *t* (defined as 1 on trials where reward occurred), and (*R*_*t*_ – *V*_*i,t*_) the prediction error. On trials where punishment occurred *R*_*t*_ = 0, which led to a decrease in value of *V*_*i*_. Model 3 also included the stimulus stickiness parameter. The final quantity controlling choice was determined by *Q*_*t*_ = *Q*^*reinf*^_*t*_ + *Q*^*stim*^_*t*_.

Model 4 took a different approach, and had three parameters: φ (phi), ρ (rho), and β (beta). Derived from the experienced-weighted attraction model (EWA) of Camerer and Ho ^55^, here it was implemented as in den Ouden *et al*. ^14^ a study in which the EWA model best described behavior best on a nearly identical human task. A key difference to the other reinforcement learning models tested in this study is that here the learning rate can decline over time, governed by a decay factor ρ (rho). The EWA model weighs the value of new information against current expectations or beliefs, accumulated from previous experience.

Learning from reinforcement is modulated by an “experience weight”, *n*_*c,t*_, which is a measure of how often the subject has chosen a stimulus (*i*.*e*. experienced the action), and is updated every time the stimulus is chosen (where *c* is choice and *t* is trial) according to the experience decay factor ρ (range 0 < ρ < 1) and can increase without bounds ^14^:

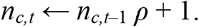

The value of a choice is updated according to the outcome, λ, and the decay factor for previous payoffs, φ (range 0 < φ < 1) ^14^

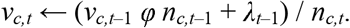

The payoff decay factor φ (phi) is related to a Rescorla–Wagner-style ^56^ learning rate α (as in Models 1-3), by α = 1 – φ. A high value of φ means that stimuli keep a high fraction of their previous value and thus learning from reinforcement is slow. When ρ is high, then “well-known” actions (with high *n*) are updated relatively little by reinforcement, by virtue of the terms involving *n*, whilst reinforcement has a proportionately larger effect on novel actions (with low *n*). For comparison to Models 1-3, when ρ = 0, the experience weight *n*, is 1, which reduces to a learning rate α controlling the influence of learning from prediction error. Choice in the EWA model is also governed by a softmax process, only here the softmax inverse temperature β was also a parameter able to vary, in contrast to Models 1-3.

## Results

### Choice of model

Behavior in all experiments was best described by reinforcement learning models incorporating parameters for stickiness, reinforcement sensitivity, and learning rates, consistent with previous work ^7, 49^. Convergence was good with most models having 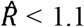 (see Methods). Model comparison metrics are shown in Supplemental Table 2. For all experiments, the winning model had separate learning rates for reward (α^*rew*^) and punishment (α^*pun*^). The reward learning rate (α^*rew*^) indexed how quickly action value representation increased following a reward prediction error (when action outcome was better than predicted). Punishment learning rate (α^*pun*^) is an assay of the speed at which action value decreased following a punishment prediction error (outcome was worse than predicted). Stickiness measures a basic perseverative tendency: whether or not an action chosen on the previous trial was repeated, irrespective of its outcome. For rats, stickiness indexed the side (or location; τ^*loc*^) of responding whereas for humans, stickiness referred to (visual) stimulus stickiness (τ^*stim*^). Reinforcement sensitivity (τ^*reinf*^) measures the degree to which the values learned through reinforcement impact on choice behavior. Reinforcement sensitivity can be viewed as a value-based inverse temperature; stickiness as a value-free inverse temperature. Low values of stickiness or reinforcement sensitivity can be thought of as two different types of exploratory behavior; low reinforcement sensitivity represents exploration away from the more highly valued choice whereas low stickiness represents exploration away from the previously chosen stimulus or location irrespective of value. The accuracy of the parameter recovery was confirmed for this modelling approach previously ^7^ and also confirmed by simulations for those parameter values estimated here in each experiment (Supplementary Table 3).

### Serotonin depletion by intraventricular 5,7 dihydroxytryptamine (5,7-DHT): rats

Results are shown in Figure 1C and Table 1. Post-mortem neurochemistry confirmed that 5,7-DHT infusions produced a near-total depletion of brain serotonin (for more details see the Methods and also Bari *et al*. 2010). The conventional analysis in the previous publication ^19^ found a decreased win-stay rate, an increased lose-shift rate and a reduced number of reversals completed in the group of depletion-operated rats (*n* = 8) compared with the group of sham-operated rats (*n* = 8). After computational modelling, we found that the depletion decreased the side (location) stickiness parameter (τ^*loc*^; MD = -0.2938 [95% HDI, -0.4635 to - 0.1134]) and the reward learning rate (α^*rew*^; MD = -0.0401 [85% HDI, -0.0757 to -0.0033]).

There was no effect of 5,7-DHT on the punishment learning rate (α^*pun*^) or reinforcement sensitivity (τ^*reinf*^) [0 ∈ 75% HDI]. The decreased lose-shift rate was retrodicted in the simulation of the computational model (Supplementary Result 1). Furthermore, because reinforcement sensitivity was also unaffected in Model 1, which did not contain the stickiness parameter, the effect of 5,7-DHT on stickiness was unlikely to be a misattribution of reinforcement sensitivity.

### Acute SSRI: rats

Results for acute citalopram administered to rats (*n* = 11 with a cross-over design for vehicle, 1mg/kg, and 10mg/kg) are shown in Figure 2 and Table 1. The conventional analysis showed the number of reversals completed was significantly lower following a low dose of 1 mg/kg SSRI compared with a high dose of 10 mg/kg SSRI ^19^. After computational modelling of the behavior, we found a single dose of 1 mg/kg citalopram in rats diminished the side (location) stickiness parameter (MD = -0.1862 [95% HDI, -0.3330 to -0.0441]), as seen following 5,7-DHT. The reward learning rate was enhanced by the 1 mg/kg dose in rats (MD = 0.2098 [95% HDI, 0.0184 to 0.3959]). There was no effect of 1 mg/kg on the punishment learning rate or reinforcement sensitivity (0 ∈ 75% HDI). A single high dose of citalopram in rats (10 mg/kg) decreased the reward learning rate (MD = -0.1489 [85% HDI, -0.2888 to -0.0009]) and enhanced reinforcement sensitivity (MD = 0.2900 [85% HDI, 0.0346 to 0.5590]). However, there was no effect of 10 mg/kg on the punishment learning rate or side (location) stickiness (0 ∈ 75% HDI). Simulation of the wining model retrodicted the significant difference in the number of reversals completed between the low-dose group and the high-dose group (Supplementary Result 1).

**Figure 2.**
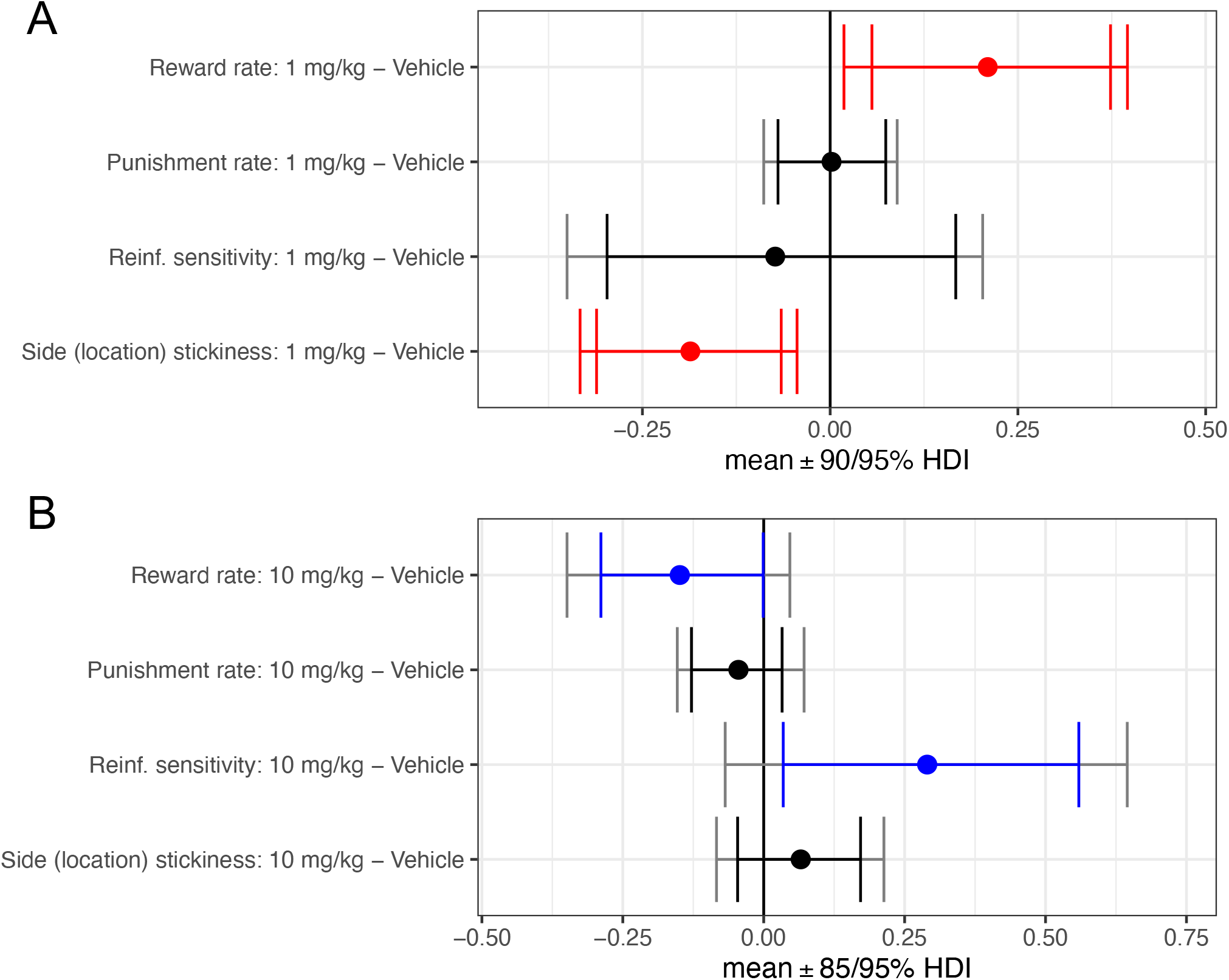
Effects of acute SSRI (citalopram) at two doses on model parameters in rats. **A**) for 1 mg/kg and **B**) for 10 mg/kg. Reinf. = reinforcement. mg/kg = milligrams per kilogram. Red signifies a difference between the parameter per-condition mean according to the Bayesian “credible interval”, 0 ∉ 95% HDI. Blue signifies a significance by the 85% HDI. The inner interval stands for the 90% HDI in A), and 85% HDI in B), while the outer interval represents the 95% HDI.

### Repeated and sub-chronic SSRI: rats

Results for ‘repeated’ 5 mg/kg citalopram administered for consecutive 7 days to rats (the Cit group; *n* = 7) compared with the vehicle group (the Veh group; *n* = 7) are shown in Figure 3A and Table 1. After 7 days, the Cit group received 10 mg/kg of citalopram twice a day for 5 consecutive days to study the longer-lasting effects of ‘sub-chronic’ dosing. Results for sub-chronic dosing are shown in Figure 3B and Table 1. The conventional analyses showed the win-stay rate increased by repeated citalopram treatment and the number of reversals was increased by sub-chronic dosing ^19^. Following computational modelling of the behavior, we found that repeated citalopram enhanced both the punishment learning rate (MD = 0.3299 [95% HDI, 0.0432 to 0.6404]) and side (location) stickiness (MD = 0.1581 [75% HDI, 0.0135 to 0.3054]). There was no effect of repeated citalopram on the reward learning rate and reinforcement sensitivity (0 ∈ 75% HDI). The sub-chronic dosing enhanced the reward learning rate (MD = 0.4769 [95% HDI, 0.2699 to 0.6780]), the punishment learning rate (MD = 0.4762 [95% HDI, 0.2172 to 0.7323]), and the side (location) stickiness (MD = 0.1676 [75% HDI, 0.0075 to 0.3414]), but decreased the reinforcement sensitivity (MD = -0.9972 [95% HDI, -1.7233 to -0.2540]). Simulation of the winning model retrodicted the significant increase of the win-stay rate for repeated citalopram compared with the vehicle, but did not show a significant increase in the number of reversals for sub-chronic dosing (Supplementary Result 1).

**Figure 3.**
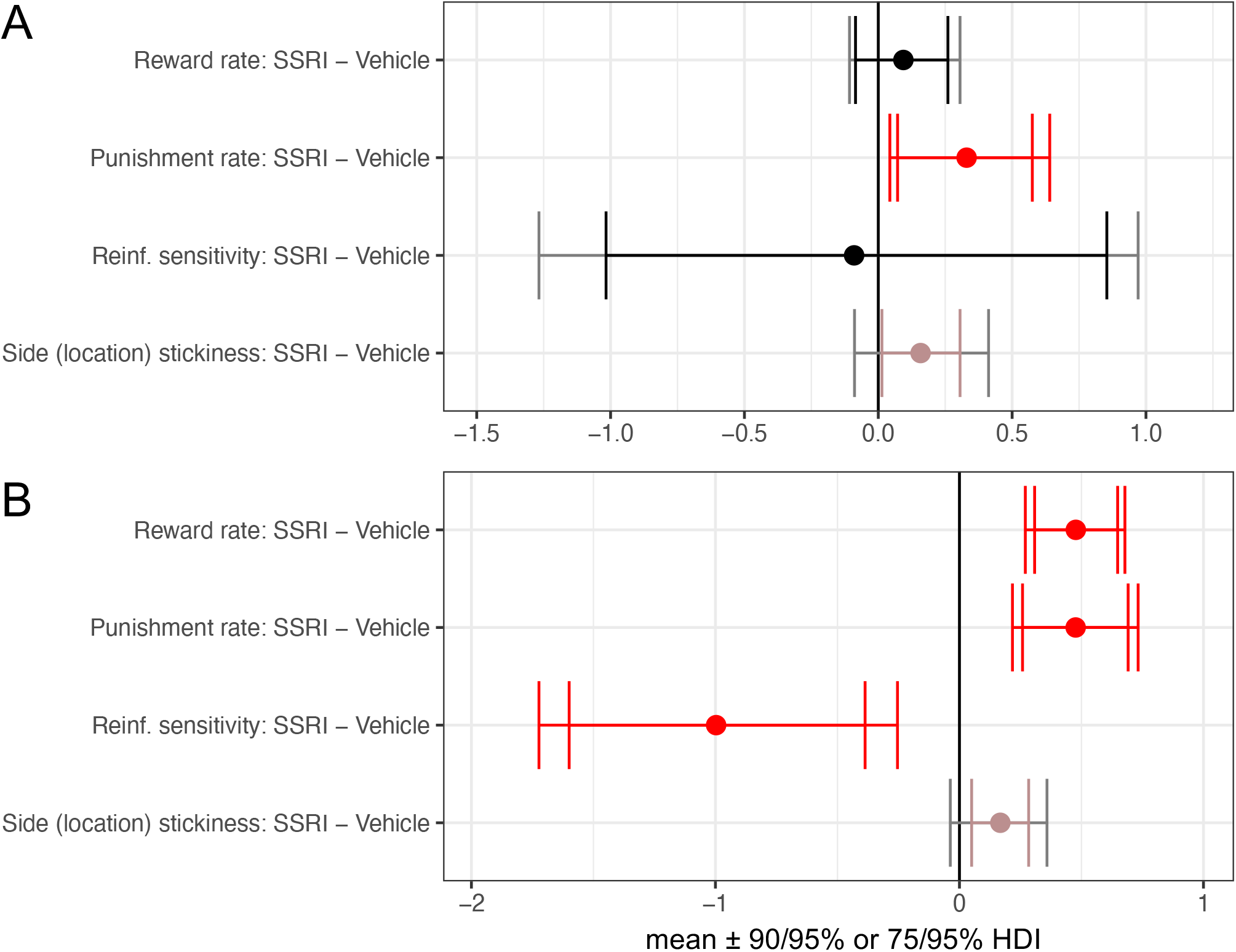
Effects of repeated and sub-chronic SSRI on model parameters in rats. **A**) for the repeated SSRI (5 mg/kg citalopram) experiment, and **B**) for the sub-chronic SSRI (10 mg/kg citalopram) experiment. Reinf. = reinforcement. Red signifies a difference between the parameter per-condition mean according to the Bayesian “credible interval”, 0 ∉ 95% HDI, and orange signifies a significance by the 75% HDI. All outer intervals represent the 95% HDI. The inner intervals represent the 75% HDI for side stickiness and the 90% HDI for the other 3 parameters.

### Acute SSRI: humans

Modelling results (*n* = 32 escitalopram, *n* = 33 placebo) are shown in Figure 4 and Table 1. The prior conventional analysis suggested that the impaired reversal learning after acute SSRI mainly resulted from an elevated lose-shift rate ^22^. After computational modelling, we found that the administration of a single 20 mg dose of escitalopram to healthy humans decreased the reward learning rate (MD = -0.2019 [95% HDI, -0.3612 to -0.0392]), stimulus stickiness (MD = -0.1841 [85% HDI, -0.3476 to -0.0045]) and reinforcement sensitivity (MD = -1.6848 [80% HDI, -3.1501 to -0.1553]), but had no effect on the punishment learning rate (0 ∈ 75% HDI). Simulation of the computational model retrodicted a significantly increased lose-shift rate (Supplementary Result 1).

**Figure 4.**
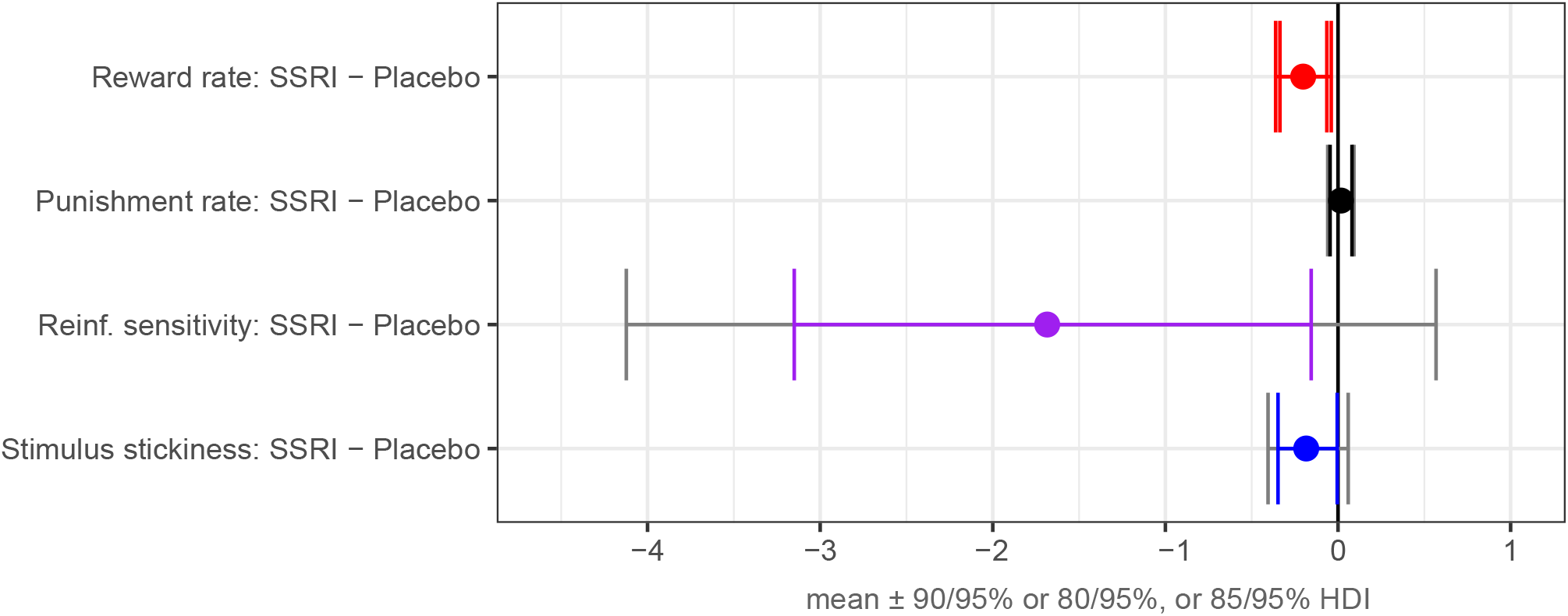
Effects of acute SSRI (20 mg escitalopram) on model parameters in humans. Stimulus stickiness was decreased following acute SSRI. Reinf. = reinforcement. Red signifies a difference between the parameter per-condition mean according to the Bayesian “credible interval”, 0 ∉ 95% HDI. Similarly, blue and purple signify the significance levels by 85% and 80% HDI’s, respectively. All outer intervals represent the 95% HDI.

### Chronic SSRI treatment in humans

As reported in our recent publication for the effect of chronic use of SSRI on behavioral flexibility by a double-blind, placebo-control, semi-randomized study ^57^, the computational modelling approach was applied to the behavioral data of the same probabilistic reversal learning task in healthy volunteers. The participants were semi-randomized into the treatment group (*n* = 32) receiving 20 mg escitalopram or the control group receiving the placebo for 3 to 5 weeks. The conventional analysis identified no significant group differences^57^. After computational modelling, we found that the chronic use of SSRI reduced reinforcement sensitivity compared to placebo (*n* = 34) in healthy volunteers (MD = -2.7673 [90% HDI, −5.2846 to −0.3959]), but had no effect on reward/punishment learning rates or stimulus stickiness (0 ∈ 75% HDI) ^57^.

### Relationship between model parameters and conventional behavioral measures

Next, we conducted correlational analyses to demonstrate how our modelling results compared with traditional metrics of PRL. There were converging effects across species involving stickiness. Results were corrected for multiple comparisons by false discovery rate (FDR) and are summarised in Supplemental Tables 5-7. The conventional measures examined for the rat experiments were win-stay (proportion of trials where the subject stayed with the same choice following a reward), lose-shift (proportion of trials where the subject shifted choice following punishment), and number of reversals completed ^19^. Win-stay and lose-shift were also examined in the human studies, as was perseveration ^18^. In the human SSRI acute experiment, stimulus stickiness was positively correlated with the win-stay rate (r = .51, p = .0066 on placebo; r = .62, p = .0005 following escitalopram) and also negatively correlated with the lose-shift rate (r = -.63, p = .0003 on placebo; r = -.78, p = 7.95 × 10^−7^ following escitalopram). In rats, side (location) stickiness was negatively correlated with the lose-shift rate following an acute 1 mg/kg dose of citalopram (r = -.89, p = .006), and positively correlated with the win-stay rate in the vehicle group with daily injections of 0.01 M phosphate-buffered saline for 7 days (r = .95, p = .0065). Side (location) stickiness was also positively correlated with the number of reversals achieved during the repeated administration (r = .89, p = .0205 following 5 mg/kg citalopram per day and r = .97, p = .0049 with the same number of daily injections of vehicle). Further correlations with other model parameters are reported in the Supplementary Tables 5-7.

### Summary of results

In rats, stickiness was decreased after 5,7-DHT and acute 1 mg/kg citalopram, whereas stickiness was increased after repeated 5 mg/kg citalopram and sub-chronic 10 mg/kg citalopram. In humans, stickiness was decreased following 20 mg escitalopram, similar to the effects of 5,7-DHT and low dose citalopram in rats. Also in cross-species alignment, the reward learning rate was decreased following 5,7-DHT and acute 10 mg/kg citalopram in rats as well as in humans following 20 mg escitalopram. The reward learning rate in rats was additionally increased following acute 1 mg/kg citalopram and sub-chronic 10mg/kg citalopram. The punishment learning rate was increased for both repeated 5 mg/kg citalopram and sub-chronic citalopram in rats only. Reinforcement sensitivity was increased following 10 mg/kg of citalopram and decreased during sub-chronic treatment in rats, agreeing with our own recent analysis of chronic escitalopram treatment in humans ^57^, although this parameter was also shown to be decreased in the present analysis following acute 20mg escitalopram in humans.

## Discussion

We have demonstrated converging effects of a range of bidirectional 5-HT manipulations across both rats and humans which bolsters its evolutionarily conserved role in behavioral flexibility and plasticity. Computational modelling of choice behavior indicated increases or decreases in choice repetition (‘stickiness’) or reinforcement learning rates depending upon manipulations intended to increase or decrease serotonin function, respectively. Stickiness, a basic tendency to persevere versus ‘explore’, was modulated in five serotonergic manipulations examined across both rats and humans. Stickiness was decreased by neurotoxic 5-HT depletion in rats and by acute 1 mg/kg SSRI in rats (citalopram) and healthy humans (20 mg escitalopram), treatments presumably reducing 5-HT signalling. By contrast, stickiness was increased following both repeated (5 mg/kg for 7 days) and sub-chronic (10 mg/kg twice a day for 5 days) dosing of SSRI in rats, treatments probably boosting 5-HT function. Learning rates were also modulated by five serotonergic manipulations across species. The reward learning rate increased the most after sub-chronic administration of the SSRI citalopram (5 mg/kg for 7 days followed by 10 mg/kg twice a day for 5 days) compared with the vehicle group. Conversely, humans given a single dose of an SSRI (20mg escitalopram), which can decrease post-synaptic serotonin signalling, and rats that received 5,7-DHT demonstrated decreased reward learning rates. This in turn parallels the reduction of reinforcement learning rates following 5,7-DHT infused directly in the marmoset amygdala or OFC to produce local 5-HT depletion ^46^. Collectively, the present and the previous results show that serotonin has common effects on latent computational mechanisms supporting flexible decision-making and plasticity in rats, marmoset monkeys and humans.

The neural substrates of PRL are relatively well understood ^46, 58, 59^ and involve interactions in particular among the orbitofrontal cortex (OFC), amygdala, and striatum. Administration of 5,7-DHT directly to either the marmoset OFC or amygdala produced changes in both stickiness and reinforcement learning rates ^46^. Marmosets that received 5,7-DHT in the OFC repeated choices to recently chosen stimuli across a longer timescale, whereas 5,7-DHT in the amygdala produced a more ephemeral tendency to repeat choices ^46^. Dietary depletion of tryptophan, serotonin’s biosynthetic precursor, in humans, also modulated stickiness and corresponding activity in frontopolar cortex during a four-choice probabilistic task ^60^.

Stickiness, the only value-free parameter in our reinforcement learning model, contributed to a core feature of complex behavior, *i*.*e*. exploration. Lower stickiness, even negative stickiness, is generally associated with more exploratory behavior. However, exploratory behavior is not a unitary construct ^61^. At one level, exploratory behavior can reflect directed information gathering, but on another level it can be mechanistic or rigid, resulting from ‘decisional noise’, producing apparently flexible behavior but, in fact, representing a fundamental performance heuristic recruited in volatile settings that evokes a primitive form of exploration. Another potential measure of exploratory behavior is reflected in reinforcement sensitivity, as a value-based parameter in our model, which can be interpreted as reflecting the balance between exploiting and exploring tendencies (low reinforcement sensitivity is sometimes referred to as ‘random exploration’) ^62^.

Whilst the effects of serotonin on reinforcement sensitivity revealed by the present analyses were ostensibly more difficult to interpret – underscoring that stickiness is a distinct mechanism – there is an intriguing parallel with a recent study. Langley *et al*. ^57^ have recently shown diminished reinforcement sensitivity in healthy humans following chronic – at least 21 days – of 20 mg escitalopram performing the same PRL task and modelled in an identical fashion – this reduction is hence the same direction as for the acute dose in humans and sub-chronic dosing in rats. Although this parallel between single and chronic dosing in humans was unexpected, it is notable that reinforcement sensitivity in rats following sub-chronic dosing was also decreased. These effects of reduced reinforcement sensitivity (value-based) may relate to what has been termed “emotional blunting” or “SSRI-induced apathy syndrome” in patients with MDD ^57, 63-65^. The reduction in inverse temperature can also be interpreted as a reduction in “maximisation” of reinforcement and this a shift in the balance between “exploitation” and “exploration” ^61^. However, it is evident that this drift to exploration is not always accompanied by reduced “stickiness”, suggesting different processes underlying choice variability.

The present analyses focusing on behavioral flexibility are relevant to current hypotheses of effects of psychedelic agents such as psilocybin and LSD and their hypothetical actions on neuronal plasticity and cognitive flexibility ^66, 67^. There are in fact intriguing parallels between the present global manipulations of serotonin and the effects of LSD on latent mechanisms underlying PRL in humans. Whilst LSD is mostly known for its 5-HT_2A_ agonist properties, it is also a 5-HT_1A_ agonist and suppresses dorsal raphe serotonin neuron activity ^68^. Indeed, LSD was recently shown to reduce stickiness during PRL performance of healthy humans ^69, 70^, which aligns with 5-HT_1A_ somatodendritic autoreceptor effects associated with the reduced stickiness shown here following acute SSRI in humans and low dose SSRI in rats. At the same time, LSD markedly increased the reinforcement learning rates for both reward and punishment ^70^, which were also increased following sub-chronic SSRI dosing in rats. The parallel with our sub-chronic SSRI results from rats with the effects of LSD on learning rates in humans agrees with the literature showing that optogenetic stimulation of 5-HT neurons in the dorsal raphe increased reinforcement learning rates^71^. Given the well-established role of the 5-HT_2A_ receptor in reversal learning, and its involvement in SSRI-related reversal improvements ^72^, a 5-HT_2A_ mechanism may well be implicated in the present data. Indeed, the 5-HT_2A_ receptor is involved in plasticity ^73, 74^ and associative learning ^75^. Furthermore, during initial learning (pre-reversal), LSD decreased reinforcement sensitivity ^70^, in line with the acute and chronic ^57^ SSRI effects in humans and sub-chronic effect in rats.

Other studies have investigated other forms of exploratory behavior, sometimes assessed with a four-choice, rather than two-choice, task as here. For example, directed exploration – where the goal is to explore uncertain options to maximise information gained – was modulated by dopamine ^76^ and attenuated in gambling disorder ^77^. *Tabula rasa* exploration (disregarding history), meanwhile, ignores all prior knowledge (*e*.*g*. choice history, reinforcement history, and estimates of uncertainty, respectively), has been associated with norepinephrine but not dopamine function ^78^ and may be enhanced in individuals with attention-deficit/hyperactivity disorder (ADHD) symptoms ^79^. Understanding distinct types of exploratory behavior and their neurochemical modulation is therefore relevant transdiagnostically. We posit that low stickiness is a fundamental form of exploration, and have shown here that serotonin modulates it; this is likely by affecting a neural network that includes the dorsomedial PFC, OFC, and amygdala ^46^.

Manifestation of high or low stickiness may bear on the neural representation of discrete states of the world. In the context of PRL, for example, one state would be “option A is mostly correct” (pre-reversal) whilst another state would be “option B is mostly correct” (post-reversal). To perform well during PRL, in this view, veridical state representations inferred by the brain are critical as are veridical probabilities of transitions between states. Indeed, the OFC is implicated in representing states ^80, 81^. One possibility, therefore, is that these results concerning stickiness collectively reflect an influence of serotonin on inferring states or state transitions. This would align with recent theorising on OCD (where stickiness is low during PRL) ^7^, which posits that the disorder can be characterised by excessive statistical uncertainty (variance, or inverse precision) about the probability of transitions between states (*e*.*g*. from the state of dirty hands to clean hands after washing), particularly those that are action-dependent ^82^. The optimal response to uncertainty about the current state would be exploratory behavior to continue gathering information ^82^. SUD (where stickiness is high) ^7^, meanwhile, may be characterised by over-encoding of state-specific rules and information ^83^. The model of state transition uncertainty can explain excessive behavioral switching (*i*.*e*. low stickiness) as well as heightened perseveration (*i*.*e*. high stickiness) and can be extended to account for other conditions including generalised anxiety disorder, autism spectrum disorder (ASD), and schizophrenia ^82^. Indeed, reversal learning deficits have been documented in ASD ^6^ and schizophrenia ^84, 85^.

Dose-dependent effects of SSRIs are key to understanding serotonin function in this cross-species analysis. Acute low- and high-dose SSRI administration lowered and increased stickiness, respectively, which likely reflected sensitive measures of opposite effects on 5-HT activity. Evidence from positron emission tomography (PET) imaging has shown that acute SSRI in humans, at the dose used here, lowers 5-HT concentrations in projection regions ^86^, although there can be considerable individual differences in this action^87^ - which may relate to the considerable variability in the reinforcement sensitivity parameter evident in Figure 4. The reduction in 5-HT levels in terminal projection areas is believed to reflect the activation of 5-HT_1A_ autoreceptors by increases in extracellular serotonin following reuptake inhibition, which in turn leads to decreased firing rates of 5-HT neurons ^42, 44^. We posit that the high acute dose of SSRI used in rats, which heightened stickiness, overcame 5-HT_1A_ autoreceptor-mediated regulation.

The dose-dependent effects on stickiness may have implications for the treatment of OCD, in particular, one of numerous conditions for which SSRIs are first-line pharmacotherapy ^38-41^. One puzzle has been why doses up to three times higher than those used in MDD are optimal for reducing symptoms of OCD ^88^. In fact, guidelines for OCD recommend titrating to the maximum approved dose ^89^, yet using these high doses in MDD does not improve efficacy and instead increases side-effects ^88^. That both the repeated 5 mg/kg SSRI and the sub-chronic 10 mg/kg treatments in rats increased stickiness in the present study may be relevant for understanding this clinical phenomenon.

## Conclusion

It is imperative to overcome the challenge of relating animal and human experiments in order to advance models of psychiatric disorder and drug development ^90-92^. Here, we have provided evidence across rats and humans that serotonin modulates fundamental components of learning important for plasticity (reinforcement learning rates) and behavioral flexibility (stickiness), bidirectionally. Stickiness, a basic perseverative tendency less commonly studied in conjunction with RL, may be a fundamental mechanism involved in choice. Moreover, we have shown a consistent role for serotonin in affecting basic tendencies to persevere or explore in comparable decision-making tasks in rats and humans. These results demonstrate that the role of serotonin in cognitive flexibility is preserved across species and are thus of evolutionary significance. In addition, this role of serotonin is of clinical relevance for neuropsychiatric disorders where SSRIs are the first line of treatment. The translational results of this study are of particular relevance for the pathophysiology and treatment of OCD and SUD, where parallel learning processes have been perturbed ^7^, and have implications for a wide range of other neuropsychiatric disorders, including depression ^8, 9^ and schizophrenia ^27, 93^.

## Acknowledgements

This research was funded by the Wellcome Trust (Grant 104631/Z/14/Z to T.W.R) and the Lundbeck Foundation (Grant R281-2018-131 to B.J.S and G.M.K). Q.L. was supported by the National Key Research and Development Program of China (Grant 2019YFA0709502), the National Natural Science Foundation of China (Grant 82272079), the Natural Science Foundation of Shanghai (Grant 20ZR1404900), the Shanghai Municipal Science and Technology Major Project (Grant 2018SHZDZX01). Most of the analyses of this study had been conducted when Q.L. was a Visiting Fellow at the Clare Hall, University of Cambridge, Cambridge, UK. J.W.K. was supported by a Gates Cambridge Scholarship and an Angharad Dodds John Bursary in Mental Health and Neuropsychiatry. R.N.C was funded by the UK Medical Research Council (MC_PC_17213). N.S. was supported by an Academic Clinical Fellowship (University of Cambridge/ Cambridgeshire and Peterborough NHS Foundation Trust). B.U.P was supported by an Angharad Dodds John Bursary in Mental Health and Neuropsychiatry and is a current employee of AstraZeneca plc.

## Author Contributions

TWR, QL and JWK made substantial contributions to the conception or design of the work; AB, NS and CL contributed substantially to the acquisition of the data; QL, JWK, JA, BUP and RNC contributed substantially to the analysis of the data; QL, JWK, GMK, BJS, RNC and TWR contributed substantially to the interpretation of data; JK and QL wrote the first draft; AB, NS, CL, GMK, JA, BUP, BJS, RNC and TWR made critical revisions.

## Reference

1. Berlin, G.S. & Hollander, E. Compulsivity, impulsivity, and the DSM-5 process. Cns Spectrums 19, 62–68 (2014).

2. Jentsch, J.D. & Taylor, J.R. Impulsivity resulting from frontostriatal dysfunction in drug abuse: implications for the control of behavior by reward-related stimuli. Psychopharmacology 146, 373–390 (1999).

3. Koob, G.F. & Volkow, N.D. Neurobiology of addiction: a neurocircuitry analysis. Lancet Psychiatry 3, 760–773 (2016).

4. Robbins, T.W., Vaghi, M.M. & Banca, P. Obsessive-Compulsive Disorder: Puzzles and Prospects. Neuron 102, 27–47 (2019).

5. Tiffany, S.T. A COGNITIVE MODEL OF DRUG URGES AND DRUG-USE BEHAVIOR – ROLE OF AUTOMATIC AND NONAUTOMATIC PROCESSES. Psychological Review 97, 147–168 (1990).

6. Brolsma, S.C.A., et al. Challenging the negative learning bias hypothesis of depression: reversal learning in a naturalistic psychiatric sample. Psychological Medicine 52, 303–313 (2022).

7. Kanen, J.W., Ersche, K.D., Fineberg, N.A., Robbins, T.W. & Cardinal, R.N. Computational modelling reveals contrasting effects on reinforcement learning and cognitive flexibility in stimulant use disorder and obsessive-compulsive disorder: remediating effects of dopaminergic D2/3 receptor agents. Psychopharmacology 236, 2337–2358 (2019).

8. Mukherjee, D., Filipowicz, A.L.S., Vo, K., Satterthwaite, T.D. & Kable, J.W. Reward and Punishment Reversal-Learning in Major Depressive Disorder. Journal of Abnormal Psychology 129, 810–823 (2020).

9. Murphy, F.C., Michael, A., Robbins, T.W. & Sahakian, B.J. Neuropsychological impairment in patients with major depressive disorder: the effects of feedback on task performance. Psychological Medicine 33, 455–467 (2003).

10. Taylor Tavares, J.V., et al. Neural basis of abnormal response to negative feedback in unmedicated mood disorders. Neuroimage 42, 1118–1126 (2008).

11. Alsio, J., et al. Serotonergic Innervations of the Orbitofrontal and Medial-prefrontal Cortices are Differentially Involved in Visual Discrimination and Reversal Learning in Rats. Cerebral Cortex 31, 1090–1105 (2021).

12. Brown, H.D., Amodeo, D.A., Sweeney, J.A. & Ragozzino, M.E. The selective serotonin reuptake inhibitor, escitalopram, enhances inhibition of prepotent responding and spatial reversal learning. Journal of Psychopharmacology 26, 1443–1455 (2012).

13. Clarke, H.F., Dalley, J.W., Crofts, H.S., Robbins, T.W. & Roberts, A.C. Cognitive inflexibility after prefrontal serotonin depletion. Science 304, 878–880 (2004).

14. den Ouden, H.E.M., et al. Dissociable Effects of Dopamine and Serotonin on Reversal Learning. Neuron 80, 1090–1100 (2013).

15. Lapiz-Bluhm, M.D.S., Soto-Pina, A.E., Hensler, J.G. & Morilak, D.A. Chronic intermittent cold stress and serotonin depletion induce deficits of reversal learning in an attentional set-shifting test in rats. Psychopharmacology 202, 329–341 (2009).

16. Matias, S., Lottem, E., Dugue, G.P. & Mainen, Z.F. Activity patterns of serotonin neurons underlying cognitive flexibility. Elife 6 (2017).

17. Kanen, J.W., et al. Effect of lysergic acid diethylamide (LSD) on reinforcement learning in humans. bioRxiv, 2020.2012.2004.412189 (2021).

18. Kanen, J.W., et al. Probabilistic reversal learning under acute tryptophan depletion in healthy humans: a conventional analysis. Journal of Psychopharmacology 34, 580–583 (2020).

19. Bari, A., et al. Serotonin Modulates Sensitivity to Reward and Negative Feedback in a Probabilistic Reversal Learning Task in Rats. Neuropsychopharmacology 35, 1290–1301 (2010).

20. Chamberlain, S.R., et al. Neurochemical modulation of response inhibition and probabilistic learning in humans. Science 311, 861–863 (2006).

21. Lawrence, A.D., Sahakian, B.J., Rogers, R.D., Hodges, J.R. & Robbins, T.W. Discrimination, reversal, and shift learning in Huntington’s disease: mechanisms of impaired response selection. Neuropsychologia 37, 1359–1374 (1999).

22. Skandali, N., et al. Dissociable effects of acute SSRI (escitalopram) on executive, learning and emotional functions in healthy humans. Neuropsychopharmacology 43, 2645–2651 (2018).

23. Daw, N.D. Trial-by-trial data analysis using computational models. Decis Making, Affect Learn Atten Perform XXIII, 1–26 (2011).

24. Sutton, R.S. & Barto, A.G. Reinforcement Learning: An Introduction, 2nd edn. The MIT Press (2012).

25. Bennett, D., Niv, Y. & Langdon, A.J. Value-free reinforcement learning: policy optimization as a minimal model of operant behavior. Current Opinion in Behavioral Sciences 41, 114–121 (2021).

26. Miller, K.J., Shenhav, A. & Ludvig, E.A. Habits Without Values. Psychological Review 126, 292–311 (2019).

27. Deserno, L. & Hauser, T.U. Beyond a Cognitive Dichotomy: Can Multiple Decision Systems Prove Useful to Distinguish Compulsive and Impulsive Symptom Dimensions? Biol Psychiatry 88, e49–e51 (2020).

28. Shahar, N., et al. Credit assignment to state-independent task representations and its relationship with model-based decision making. Proceedings of the National Academy of Sciences of the United States of America 116, 15871–15876 (2019).

29. Balleine, B.W. & O’Doherty, J.P. Human and Rodent Homologies in Action Control: Corticostriatal Determinants of Goal-Directed and Habitual Action. Neuropsychopharmacology 35, 48–69 (2010).

30. Cardinal, R.N., Parkinson, J.A., Hall, J. & Everitt, B.J. Emotion and motivation: the role of the amygdala, ventral striatum, and prefrontal cortex. Neuroscience and Biobehavioral Reviews 26, 321–352 (2002).

31. Ersche, K.D., et al. Carrots and sticks fail to change behavior in cocaine addiction. Science 352, 1468–1471 (2016).

32. Gillan, C.M., et al. Disruption in the Balance Between Goal-Directed Behavior and Habit Learning in Obsessive-Compulsive Disorder. American Journal of Psychiatry 168, 718–726 (2011).

33. Voon, V., et al. Disorders of compulsivity: a common bias towards learning habits. Molecular Psychiatry 20, 345–352 (2015).

34. Ohmura, Y., et al. Disruption of model-based decision making by silencing of serotonin neurons in the dorsal raphe nucleus. Current Biology 31, 2446-+ (2021).

35. Worbe, Y., et al. Valence-dependent influence of serotonin depletion on model-based choice strategy. Molecular Psychiatry 21, 624–629 (2016).

36. Worbe, Y., Savulich, G., de Wit, S., Fernandez-Egea, E. & Robbins, T.W. Tryptophan Depletion Promotes Habitual over Goal-Directed Control of Appetitive Responding in Humans. Int J Neuropsychopharmacol 18, pyv013 (2015).

37. Bjorklund, A., Baumgarten, H.G. & Rensch, A. 5,7-DIHYDROXYTRYPTAMINE - IMPROVEMENT OF ITS SELECTIVITY FOR SEROTONIN NEURONS IN CNS BY PRETREATMENT WITH DESIPRAMINE. Journal of Neurochemistry 24, 833–835 (1975).

38. Gelenberg AJ, F.M., Markowitz JC, et al. Practice guideline for the treatment of patients with major depressive disorder. American Psychiatric Association (2010).

39. Baldwin, D.S., et al. Evidence-based pharmacological treatment of anxiety disorders, post-traumatic stress disorder and obsessive-compulsive disorder: A revision of the 2005 guidelines from the British Association for Psychopharmacology. Journal of Psychopharmacology 28, 403–439 (2014).

40. Benedek DM, F.M., Zatzick D, Ursano RJ. Practice guideline for the treatment of patients with acute stress disorder and posttraumatic stress disorder. Am Psychiatr Assoc (2009).

41. Fineberg NA, D.L., Reid J, et al. Management and treatment of OCD. In: Geddes JR, Andreasen NC, Goodwin GM (eds) New Oxford Textbook of Psychiatry, 3rd edn. Oxford University Press, Oxford, England (2020).

42. Richardson-Jones, J.W., et al. 5-HT1A Autoreceptor Levels Determine Vulnerability to Stress and Response to Antidepressants. Neuron 65, 40–52 (2010).

43. Nord, M., Finnema, S.J., Halldin, C. & Farde, L. Effect of a single dose of escitalopram on serotonin concentration in the non-human and human primate brain. International Journal of Neuropsychopharmacology 16, 1577–1586 (2013).

44. Fischer, A.G., Jocham, G. & Ullsperger, M. Dual serotonergic signals: a key to understanding paradoxical effects? Trends Cogn Sci (2014).

45. Taylor, M.J., Freemantle, N., Geddes, J.R. & Bhagwagar, Z. Early onset of selective serotonin reuptake inhibitor antidepressant action - Systematic review and meta-analysis. Archives of General Psychiatry 63, 1217–1223 (2006).

46. Rygula, R., et al. Role of Central Serotonin in Anticipation of Rewarding and Punishing Outcomes: Effects of Selective Amygdala or Orbitofrontal 5-HT Depletion. Cerebral Cortex 25, 3064–3076 (2015).

47. Cardinal, R.N. & Aitken, M.R.F. Whisker: A client-server high-performance multimedia research control system. Behavior Research Methods 42, 1059–1071 (2010).

48. Rao, N. The clinical pharmacokinetics of escitalopram. Clinical Pharmacokinetics 46, 281–290 (2007).

49. Alsio, J., et al. Dopamine D2-like receptor stimulation blocks negative feedback in visual and spatial reversal learning in the rat: behavioural and computational evidence. Psychopharmacology 236, 2307–2323 (2019).

50. Carpenter, B., et al. Stan: A Probabilistic Programming Language. Journal of Statistical Software 76, 1–29 (2017).

51. Brooks, S.P. & Gelman, A. General methods for monitoring convergence of iterative simulations. Journal of Computational and Graphical Statistics 7, 434–455 (1998).

52. Gelman, A., Hill, J. & Yajima, M. Why We (Usually) Don’t Have to Worry About Multiple Comparisons. Journal of Research on Educational Effectiveness 5, 189–211 (2012).

53. Gronau, Q.F., et al. A tutorial on bridge sampling. Journal of Mathematical Psychology 81, 80–97 (2017).

54. Gronau, Q.F., Singmann, H. & Wagenmakers, E.-J. bridgesampling: An R Package for Estimating Normalizing Constants. Journal of Statistical Software 92, 1–29 (2020).

55. Camerer, C. & Ho, T.H. Experience-weighted attraction learning in normal form games. Econometrica 67, 827–874 (1999).

56. Rescorla, R.A. & Wagner, A.R. A theory of classical conditioning: Variations in the effectiveness of reinforcement and nonreinforcement. In: Black AH, Prokasy WF (eds) Classical Conditioning II Current Research and Theory. Appleton-Century-Crofts (1972).

57. Langley, C., et al. Chronic escitalopram in healthy volunteers has specific effects on reinforcement sensitivity: a double-blind, placebo-controlled semi-randomised study. Neuropsychopharmacology (2023).

58. O’Doherty, J., Kringelbach, M.L., Rolls, E.T., Hornak, J. & Andrews, C. Abstract reward and punishment representations in the human orbitofrontal cortex. Nature Neuroscience 4, 95–102 (2001).

59. Cools, R., Clark, L., Owen, A.M. & Robbins, T.W. Defining the Neural Mechanisms of Probabilistic Reversal Learning Using Event-Related Functional Magnetic Resonance Imaging. The Journal of Neuroscience 22, 4563 (2002).

60. Seymour, B., Daw, N.D., Roiser, J.P., Dayan, P. & Dolan, R. Serotonin Selectively Modulates Reward Value in Human Decision-Making. Journal of Neuroscience 32, 5833–5842 (2012).

61. Wilson, R.C., Geana, A., White, J.M., Ludvig, E.A. & Cohen, J.D. Humans Use Directed and Random Exploration to Solve the Explore-Exploit Dilemma. Journal of Experimental Psychology-General 143, 2074–2081 (2014).

62. Gershman, S.J. Deconstructing the human algorithms for exploration. Cognition 173, 34–42 (2018).

63. Barnhart, W.J., Makela, E.H. & Latocha, M.J. SSRI-induced apathy syndrome: a clinical review. J Psychiatr Pract 10, 196–199 (2004).

64. Price, J., Cole, V. & Goodwin, G.M. Emotional side-effects of selective serotonin reuptake inhibitors: qualitative study. Br J Psychiatry 195, 211–217 (2009).

65. Marazziti, D., et al. Emotional Blunting, Cognitive Impairment, Bone Fractures, and Bleeding as Possible Side Effects of Long-Term Use of SSRIs. Clin Neuropsychiatry 16, 75–85 (2019).

66. Carhart-Harris, R.L. & Friston, K.J. REBUS and the Anarchic Brain: Toward a Unified Model of the Brain Action of Psychedelics. Pharmacol Rev 71, 316–344 (2019).

67. Vollenweider, F.X. & Preller, K.H. Psychedelic drugs: neurobiology and potential for treatment of psychiatric disorders. Nature Reviews Neuroscience 21, 611–624 (2020).

68. Nichols, D.E. Hallucinogens. Pharmacology & Therapeutics 101, 131–181 (2004).

69. Kanen, J.W., et al. Probabilistic reversal learning under acute tryptophan depletion in healthy humans: a conventional analysis. J Psychopharmacol 34, 580–583 (2020).

70. Kanen, J.W., et al. Effect of lysergic acid diethylamide (LSD) on reinforcement learning in humans. Psychol Med, 1–12 (2022).

71. Iigaya, K., Fonseca, M.S., Murakami, M., Mainen, Z.F. & Dayan, P. An effect of serotonergic stimulation on learning rates for rewards apparent after long intertrial intervals. Nature Communications 9, 2477 (2018).

72. Furr, A., Lapiz-Bluhm, M.D. & Morilak, D.A. 5-HT2A receptors in the orbitofrontal cortex facilitate reversal learning and contribute to the beneficial cognitive effects of chronic citalopram treatment in rats. The international journal of neuropsychopharmacology 15, 1295–1305 (2012).

73. Barre, A., et al. Presynaptic serotonin 2A receptors modulate thalamocortical plasticity and associative learning. Proc Natl Acad Sci U S A 113, E1382–1391 (2016).

74. Vaidya, V.A., Marek, G.J., Aghajanian, G.K. & Duman, R.S. 5-HT2A receptor-mediated regulation of brain-derived neurotrophic factor mRNA in the hippocampus and the neocortex. J Neurosci 17, 2785–2795 (1997).

75. Harvey, J.A. Role of the serotonin 5-HT(2A) receptor in learning. Learn Mem 10, 355–362 (2003).

76. Chakroun, K., Mathar, D., Wiehler, A., Ganzer, F. & Peters, J. Dopaminergic modulation of the exploration/exploitation trade-off in human decision-making. Elife 9 (2020).

77. Wiehler, A., Chakroun, K. & Peters, J. Attenuated Directed Exploration during Reinforcement Learning in Gambling Disorder. Journal of Neuroscience 41, 2512–2522 (2021).

78. Dubois, M., et al. Human complex exploration strategies are enriched by noradrenaline-modulated heuristics. Elife 10 (2021).

79. Dubois M, B.A., Moses-Payne M, et al. Tabula-rasa exploration decreases during youth and is linked to ADHD symptoms. bioRxiv (2020).

80. Schuck, N.W., Cai, M.B., Wilson, R.C. & Niv, Y. Human Orbitofrontal Cortex Represents a Cognitive Map of State Space. Neuron 91, 1402–1412 (2016).

81. Wilson, R.C., Takahashi, Y.K., Schoenbaum, G. & Niv, Y. Orbitofrontal Cortex as a Cognitive Map of Task Space. Neuron 81, 267–279 (2014).

82. Fradkin, I., Adams, R.A., Parr, T., Roiser, J.P. & Huppert, J.D. Searching for an Anchor in an Unpredictable World: A Computational Model of Obsessive Compulsive Disorder. Psychological Review 127, 672–699 (2020).

83. Mueller, L.E., Sharpe, M.J., Stalnaker, T.A., Wikenheiser, A.M. & Schoenbaump, G. Prior Cocaine Use Alters the Normal Evolution of Information Coding in Striatal Ensembles during Value-Guided Decision-Making. Journal of Neuroscience 41, 342–353 (2021).

84. Deserno L, B.R., Mathys C, et al. Volatility Estimates Increase Choice Switchingand Relate to Prefrontal Activity in Schizophrenia. Biol Psychiatry Cogn Neurosci Neuroimaging (2020).

85. Waltz, J.A. & Gold, J.M. Probabilistic reversal learning impairments in schizophrenia: Further evidence of orbitofrontal dysfunction. Schizophrenia Research 93, 296–303 (2007).

86. Nord, M., Finnema, S.J., Halldin, C. & Farde, L. Effect of a single dose of escitalopram on serotonin concentration in the non-human and human primate brain. Int J Neuropsychopharmacol 16, 1577–1586 (2013).

87. da Cunha-Bang, S., et al. Measuring endogenous changes in serotonergic neurotransmission with [11C]Cimbi-36 positron emission tomography in humans. Translational Psychiatry 9, 134 (2019).

88. Derksen, M., Feenstra, M., Willuhn, I. & Denys, D. The serotonergic system in obsessive-compulsive disorder. in Handbook of the Behavioral Neurobiology of Serotonin, 2nd Edition (ed. C.P. Muller & K.A. Cunningham) 865–891 (2020).

89. Koran, L.M., Hanna, G.L., Hollander, E., Nestadt, G. & Simpson, H.B. Practice guideline for the treatment of patients with obsessive-compulsive disorder. American Journal of Psychiatry 164, 5–53 (2007).

90. Benzina, N., N’Diaye, K., Pelissolo, A., Mallet, L. & Burguiere, E. A cross-species assessment of behavioral flexibility in compulsive disorders. Communications Biology 4 (2021).

91. Robbins, T.W. & Cardinal, R.N. Computational psychopharmacology: a translational and pragmatic approach. Psychopharmacology 236, 2295–2305 (2019).

92. Robble, M.A., et al. Concordant neurophysiological signatures of cognitive control in humans and rats. Neuropsychopharmacology 46, 1252–1262 (2021).

93. Frith, C.D. & Done, D.J. STEREOTYPED RESPONDING BY SCHIZOPHRENIC-PATIENTS ON A 2-CHOICE GUESSING TASK. Psychological Medicine 13, 779–786 (1983).

