## Supplementary Materials for "Common roles for serotonin in rats and humans for computations underlying flexible decision-making"

Qiang Luo, *et al*.

**Supplemental Result 1: Simulation Results**

*Simulation for serotonin depletion in rats*. We simulated the behaviour of 40 virtual rats in each group for 200 trials using the wining model (i.e. Model 2). We found that 5-HT depletion via 5,7-DHT significantly reduced the number of reversals completed (W = 1292.5, p = 5.6×10^-7^, by the Wilcoxon Rank Sum test for group difference in mean), reduced the win-stay rate (W=1347.5, p=1.4×10^-7^) and increased the lose-shift rate (W=415.5, p=2.2×10^-4^). These findings were consistent with the results from the conventional analysis reported in Bari et al. (2010).

*Simulation for acute SSRI in rats.* Again, we simulated 40 rats for each of the three groups for 200 trials using the winning model (i.e. Model 2). Compared with the low-dose (1 mg/kg) group, the high-dose (10 mg/kg) group completed more reversals (W=581.5, p=0.0333) and reduced the lose-shift rate (W=1284, p<10^-9^). These findings were consistent with the results from the conventional analysis reported in Bari *et al*. (2010).

*Simulation for repeated and sub-chronic SSRI in rats.* We simulated 40 rats for each of the 4 groups (repeated 5 mg/kg citalopram vs. vehicle, and sub-chronic 10 mg/kg citalopram vs. vehicle) for 200 trials using the winning model (i.e. Model 2). Compared with the vehicle group, the repeated 5 mg/kg citalopram group completed more reversals (W=412.5, p=0.0001) and had an enhanced win-stay rate (W=292, p=1.04×10^-6^). Using conventional analyses, Bari *et al*. (2010) had reported that the win-stay rate was significantly enhanced in the citalopram group compared with the vehicle group, while the increase in the number of reversals was only at a trend level. For the sub-chronic dosing group, the conventional analysis of the experimental data showed a significant increase in the number of reversals completed but no significant change in either the win-stay or the lose-shift rates when compared with the vehicle group (Bari *et al*. 2010). However, in the simulation, we found no significant change in these conventional measures (p>0.05).

*Simulation for acute SSRI in humans.* We simulated 40 virtual human participants for 80 trials (the reversal happened on the 41^st^ trial), and found a lower win-stay rate (W= 4916, p=4.32×10^-9^) and higher lose-shift rate (W=1492, p= 5.5×10^-9^) in the SSRI group compared with the placebo group. Consistent with the data reported originally by Skandali et al. (2018), in the simulation, the number of errors increased after SSRI administration (W=2556, p=0.0275) and this was particularly true during the acquisition phase of the task (W=2556, p=0.0271).

**Supplemental Figure 1.** Group comparison of conventionally behavioural measures averaged over 7 sessions for the depletion experiment in rats.

Upper row shows the data before outlier removal (the average number of reversals over 7 sessions less than 1). Two outliers were removed from the rats with depletion. Lower row shows the data after outlier removal.

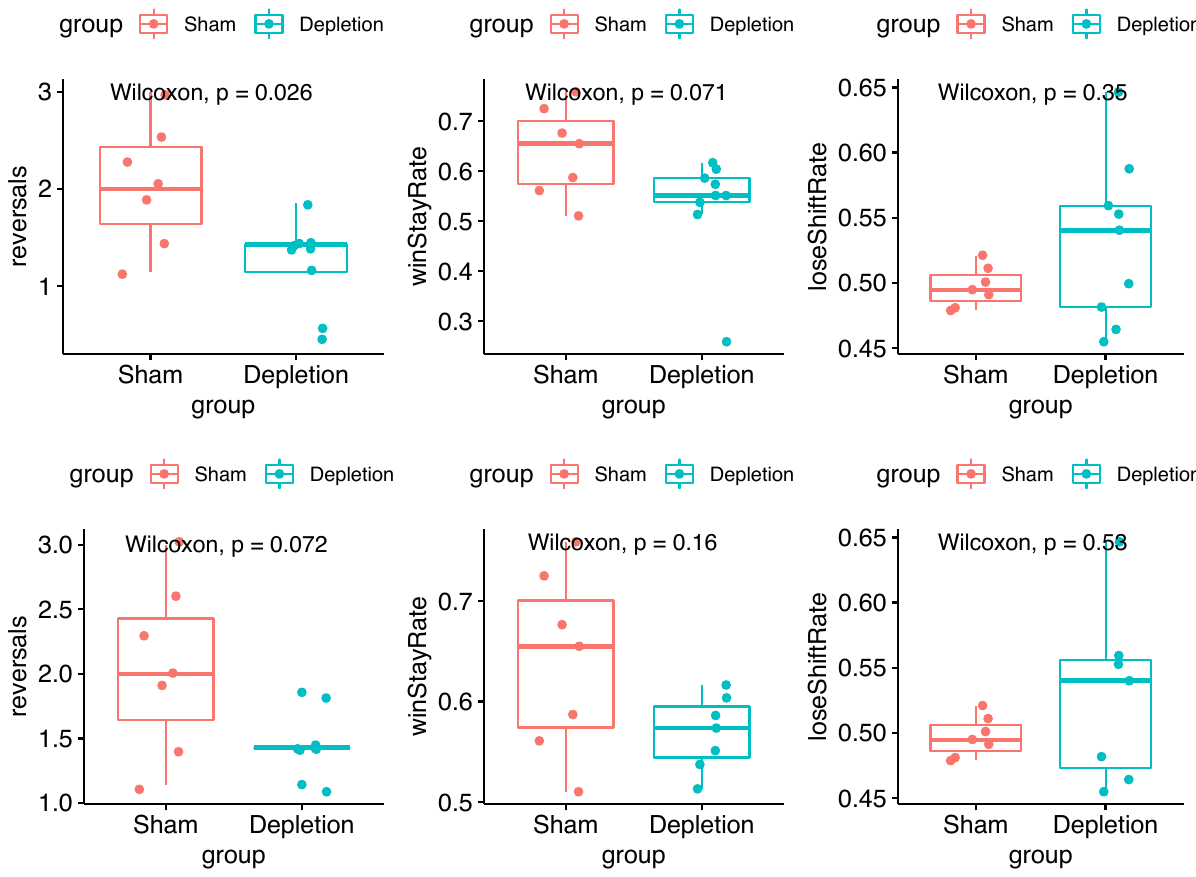

**Supplemental Table 1**. Prior distributions for model parameters

|  | **Models using each parameter** | **Prior** | **Reference** |
| --- | --- | --- | --- |
| **Model parameters** |  |  |  |
| reward learning rate, *α^rew^* | 1, 2 | Beta(1.2, 1.2) | den Ouden et al. (2013) |
| punishment learning rate, *α^pun^* | 1, 2 | Beta(1.2, 1.2) | den Ouden et al. (2013) |
| combined reward/punishment learning rate, *α^reinf^* | 3 | Beta(1.2, 1.2) | den Ouden et al. (2013) |
| reinforcement sensitivity, *τ^reinf^* | 1, 2, 3 | Gamma(*α*=4.82*, β*=0.88) | Gershman (2016) |
| stimulus stickness, *τ^stim^*  or side (location) stickness, *τ^loc^* | 2, 3 | Normal(0, 1) | Christakou et al. (2013) |
| experience decay factor, *ρ* | 4 | Beta(1.2, 1.2) | den Ouden et al. (2013) |
| decay factor for previous payoffs, *φ* | 4 | Beta(1.2, 1.2) | den Ouden et al. (2013) |
| softmax inverse temperature, *β* | 4 [note that *β =* 1 in all other models] | Gamma(*α*=4.82*, β*=0.88) | Gershman (2016) |
| **Intersubject variability in parameters** |  |  | – |
| Intersubject standard deviations for *α^rew^*, *α^pun^*, *α^reinf^, ρ , φ* | As above | Half-normal: Normal(0, 0.05) constrained to ≥0 | Kanen et al. (2019) |
| Intersubject standard deviations for *τ^reinf^, β* | As above | Half-normal: Normal(0, 1) constrained to ≥0 | Kanen et al. (2019) |

*rew* reward, *pun* punishment, *reinf* reinforcement, *stim* stimulus, *loc* location

**Supplemental Table 2.** Statistics for model comparisons.

| **Experiment** | **Rank** | **Name** | **max_rhat** | **log marginal likelihood** | **log posterior P(model)** |
| --- | --- | --- | --- | --- | --- |
| 5,7-DHT  (rats) | 2 | Model 1 | 1.5 | -14470.99 | -119.0791 |
|  | **1** | **Model 2** | 1.0 | -14351.91 | 0.0000 |
|  | 4 | Model 3 | 1.0 | -14592.29 | -240.3807 |
|  | 3 | Model 4 | 1.0 | -14508.16 | -156.2528 |
| Acute SSRI  (rats) | 4 | Model 1 | 1.0 | -3711.99 | -59.1169 |
|  | **1** | **Model 2** | 1.0 | -3652.87 | -0.0006 |
|  | 2 | Model 3 | 1.0 | -3660.24 | -7.3654 |
|  | 3 | Model 4 | 1.0 | -3709.78 | -56.9105 |
| Chronic SSRI  (rats) | 2 | Model 1 | 1.0 | -12391.85 | -235.9039 |
|  | **1** | **Model 2** | 1.0 | -12155.95 | 0.0000 |
|  | 4 | Model 3 | 1.0 | -12204.65 | -48.6963 |
|  | 3 | Model 4 | 1.0 | -12398.79 | -242.8457 |
| Sub-chronic SSRI  (rats) | 3 | Model 1 | 1.0 | -8735.83 | -59.8212 |
|  | **1** | **Model 2** | 1.0 | -8676.01 | -0.0005 |
|  | 2 | Model 3 | 1.0 | -8683.63 | -7.6163 |
|  | 4 | Model 4 | 1.7 | -8760.39 | -84.3796 |
| Acute SSRI  (humans) | 2 | Model 1 | 1.0 | -1771.95 | -7.6840 |
|  | **1** | **Model 2** | 1.0 | -1764.26 | -0.0005 |
|  | 4 | Model 3 | 1.0 | -1821.33 | -57.0660 |
|  | 3 | Model 4 | 1.0 | -1783.84 | -19.5782 |

The model ranked 1st was the winning model. Model names and parameters correspond to Table 2. Log marginal likelihood and log posterior P (model) are comparison metrics used to determine the best model. A numerically larger (less negative) log marginal likelihood is better. *rew* reward, *pun* punishment, *reinf* reinforcement, *loc* location

**Supplemental Table 3.** Simulations for parameter recovery of the parameter values estimated for each group.

| Group | Reward learning rate, *α^rew^* | Punishment learning rate, *α^pun^* | Reinforcement sensitivity, *τ^reinf^* | Stimulus stickness, *τ^stim^*  or side (location) stickness, *τ^loc^* |
| --- | --- | --- | --- | --- |
| 5,7-DHT | 97% | 97% | 100% | 97% |
| Sham | 97% | 97% | 90% | 97% |
| SSRI 1mg/kg | 97% | 97% | 97% | 97% |
| 10 mg/kg SSRI | 90% | 97% | 93% | 90% |
| Vehicle group | 97% | 90% | 97% | 93% |
| Chronic 5 mg/kg | 97% | 93% | 100% | 100% |
| Chronic sham | 97% | 97% | 93% | 83% |
| Sub-chronic 10 mg/kg | 97% | 93% | 93% | 93% |
| sub-chronic sham | 97% | 97% | 97% | 93% |
| SSRI in humans | 90% | 90% | 93% | 97% |
| Placebo in humans | 97% | 97% | 97% | 100% |

The estimated parameters were used to simulate the winning model for 100 virtual subjects. The reversal occurred on the 41^st^, of 80 trials. One choice resulted in reward on 80% of trials and the other choice resulted in reward on 20% of trials. The parameters were then fitted from these simulated data. If the 95% HDI of an estimation included the corresponding true value, then we counted this parameter recovery as a success. The models were simulated 30 times for each group. The success rates of the parameter recovery were reported.

**Supplemental Table 4.** Mean estimations of the model parameters given by Model 2.

| **Groups** | Reward learning rate, *α^rew^* | | Punishment learning rate, *α^pun^* | | Reinforcement sensitivity, *τ^reinf^* | | Stimulus stickness, *τ^stim^*  or side (location) stickness, *τ^loc^* | |
| --- | --- | --- | --- | --- | --- | --- | --- | --- |
|  | Mean  (sd) | 95% HDI | Mean  (sd) | 95% HDI | Mean  (sd) | 95% HDI | Mean  (sd) | 95% HDI |
| Sham-rats | 0.09  (0.02) | [0.04, 0.13] | 0.02  (0.01) | [1.03×10^-4^, 0.04] | 3.07  (0.41) | [2.22, 3.85] | 0.18  (0.07) | [0.05, 0.31] |
| 5,7-DHT-rats | 0.05  (0.02) | [0.01, 0.08] | 4.70×10^-3^  (0.01) | [3.48×10^-6^, 0.01] | 3.13  (0.38) | [2.39, 3.89] | -0.12  (0.06) | [-0.24, -0.01] |
| Vehicle-rats | 0.67  (0.08) | [0.51, 0.83] | 0.95  (0.04) | [0.88, 1.00] | 2.23  (0.39) | [1.48, 2.99] | 0.22  (0.08) | [0.06, 0.37] |
| 1mg/kg-rats | 0.88  (0.07) | [0.76, 1.00] | 0.95  (0.04) | [0.88, 1.00] | 2.15  (0.38) | [1.46, 2.93] | 0.03  (0.08) | [-0.12, 0.20] |
| 10mg/kg-rats | 0.52  (0.08) | [0.38, 0.67] | 0.91  (0.05) | [0.81, 1.00] | 2.52  (0.41) | [1.73, 3.30] | 0.29  (0.08) | [0.12, 0.45] |
| Placebo-human | 0.59  (0.07) | [0.46, 0.72] | 0.20  (0.03) | [0.15, 0.25] | 7.55  (0.88) | [5.91, 9.35] | 0.36  (0.09) | [0.18, 0.53] |
| SSRI-human | 0.39  (0.05) | [0.29, 0.49] | 0.21  (0.03) | [0.15, 0.27] | 5.86  (0.82) | [4.21, 7.46] | 0.18  (0.08) | [0.02, 0.34] |
| Vehicle-rats | 0.26  (0.07) | [0.11, 0.40] | 0.27  (0.11) | [0.06, 0.48] | 2.30  (0.41) | [1.50, 3.11] | 0.11  (0.09) | [-0.08, 0.28] |
| Repeated-rats | 0.35  (0.07) | [0.21, 0.51] | 0.60  (0.11) | [0.38, 0.81] | 2.21  (0.41) | [1.40, 2.99] | 0.27  (0.09) | [0.10, 0.44] |
| Vehicle-rats | 0.25  (0.06) | [0.13, 0.38] | 0.38  (0.10) | [0.19, 0.57] | 2.36  (0.30) | [1.78, 2.97] | -0.03  (0.07) | [-0.17, 0.11] |
| Sub-chronic-rats | 0.73  (0.08) | [0.56, 0.88] | 0.86  (0.09) | [0.69, 1.00] | 1.36  (0.23) | [0.92, 1.83] | 0.14  (0.07) | [0.01, 0.28] |

**Supplemental Table 5.** Correlations between model parameters and conventional measures in rats: SSRI experiments.

|  |  | **Side (Location) Stickiness, *τ^loc^*** | **Reward Learning Rate, *α^rew^*** | **Punishment Learning Rate, *α^pun^*** | **Reinforcement Sensitivity, *τ^reinf^*** |
| --- | --- | --- | --- | --- | --- |
| **Win-Stay** | **Vehicle** | -- | -- | r = .87  p = .007 | r = .94  p = .001 |
|  | **1 mg / kg Citalopram** | -- | -- | r = .94  p = .001 | r = .84  p = .009 |
|  | **10 mg / kg**  **Citalopram** | -- | -- | r = .87  p = .007 | -- |
| **Lose-Shift** | **Vehicle** | -- | -- | -- | -- |
|  | **1 mg / kg Citalopram** | r = -.89  p = .006 | -- | -- | r = .85  p = .009 |
|  | **10 mg / kg Citalopram** | -- | -- | -- | -- |
| **Reversals** | **Vehicle** | -- | -- | -- | -- |
|  | **1 mg / kg Citalopram** | -- | -- | -- | -- |
|  | **10 mg / kg Citalopram** | -- | -- | -- | -- |

Statistics reported for correlations significant at p < .05 after correction for multiple comparisons. -- = not significant. *rew* reward, *pun* punishment, *reinf* reinforcement, *loc* location

**Supplemental Table 6.** Correlations between model parameters and conventional measures in rats: repeated 5mg/kg citalopram experiment.

|  |  | **Side (location) Stickiness, *τ^stim^*** | **Reward Learning Rate, *α^rew^*** | **Punishment Learning Rate, *α^pun^*** | **Reinforcement Sensitivity, *τ^reinf^*** |
| --- | --- | --- | --- | --- | --- |
| **Win-Stay** | **Vehicle** | r = .95  p = .0065 | r=.94  p=.0065 | -- | r=-.96  p = .0059 |
|  | **Repeated** | -- | r=.96  p=.0059 | r=.89  p=.0205 | -- |
| **Lose-Shift** | **Vehicle** | -- | -- | -- | -- |
|  | **Repeated** | -- | -- | -- | -- |
| **Reversals** | **Vehicle** | r = .97  p = .0049 | r = .09  p = .0181 | -- | r = -.92  p = .0150 |
|  | **Repeated** | r=.89  p=.0205 | -- | -- | -- |

Statistics reported for correlations significant at p < .05 after correction for multiple comparisons. -- = not significant. *rew* reward, *pun* punishment, *reinf* reinforcement, *loc* location

**Supplemental Table 7.** Correlations between model parameters and conventional measures in humans: SSRI experiment.

|  |  | **Stimulus Stickiness, *τ^stim^*** | **Reward Learning Rate, *α^rew^*** | **Punishment Learning Rate, *α^pun^*** | **Reinforcement Sensitivity, *τ^reinf^*** |
| --- | --- | --- | --- | --- | --- |
| **Win-Stay** | **Placebo** | r = .51;  p = .0066 | -- | r = .44;  p = .0217 | r = .90;  p = 6.05 × 10^-12^ |
|  | **Escitalopram** | r = .62;  p = .0005 | -- | -- | r = .93;  p = 2.99 × 10^-13^ |
| **Lose-Shift** | **Placebo** | r = -.63;  p = .0003 | -- | -- | r = -.91;  p = 5.53 × 10^-12^ |
|  | **Escitalopram** | r = -.78;  p = 7.95 × 10^-7^ | -- | -- | r = -.85;  p = 3.51 × 10^-9^ |
| **Perseveration** | **Placebo** | -- | -- | -- | r = .50;  p = .0078 |
|  | **Escitalopram** | -- | -- | -- | r = .45;  p = .0217 |

Statistics reported for correlations significant at p < .05 after correction for multiple comparisons. -- = not significant. *reinf* reinforcement, *stim* stimulus
